# Short-term effects of experimental artificial light at night on nocturnal vocal activity in a temperate bird community

**DOI:** 10.64898/2026.09.16.751950

**Authors:** Kinga Buda, Jakub Buda, Michał Budka

## Abstract

Artificial light at night (ALAN) can alter the daily timing of birds activity, but the immediate effects of brief nocturnal illumination on vocal behaviour remain poorly understood. We experimentally examined whether short-term exposure to a local artificial light source affected nocturnal vocalisation rates and whether responses differed among species and between stages of the breeding season. Experiments were conducted at ten sampling points in western Poland, during nautical night in the early and late stages of the breeding season. Bird vocalisations were recorded continuously for 30 min before illumination, for 38 min during gradual and full illumination, and for 30 min after the lights were switched off. We recorded 28,839 vocalisations from 13 species. Of the five species analysed formally, only the Eurasian coot showed a significant illumination effect, with a higher vocalisation rate after illumination than before or during illumination. The sedge warbler showed higher estimated rates during and after illumination than before illumination, but the overall phase effect was not significant after false-discovery-rate correction. No phase effects were detected in the common nightingale, common quail, or Savi’s warbler. Descriptive patterns show increased vocal activity during or after illumination in the Eurasian skylark, common cuckoo, and corn crake, but limited spatial replication prevented formal testing. Overall, brief illumination did not produce a clear and consistent response across the recorded species. Short-term ALAN exposure may affect nocturnal vocal activity with some delay, but responses appear to be strongly species-specific. The observed patterns identify candidate species for further controlled experimental study.

## INTRODUCTION

Light is a major environmental cue that entrains avian circadian rhythms (Cassone, 2014). Artificial light at night (ALAN) alters natural light-dark cycles and affects avian physiology and behaviour, including sleep, metabolism, reproductive timing, foraging, and daily activity (Diaz-Palma et al., 2025). ALAN is commonly associated with earlier dawn vocalisation and later evening vocal activity (Da Silva & Kempenaers, 2017; Dominoni, 2015), although the magnitude and direction of these responses, as well as their potential fitness consequences, vary among species and environmental contexts (Wang et al., 2021). Recent global evidence indicates that light pollution can substantially extend the daily period of avian vocal activity but also confirms considerable interspecific variation in this response (Pease & Gilbert, 2025). Most research has examined the onset of dawn activity and the cessation of evening activity, whereas immediate responses to discrete ALAN exposure during the night remain less well understood (Da Silva et al., 2016). Nocturnal vocalisation by otherwise diurnal birds is taxonomically widespread and varies with both natural and artificial illumination (La, 2012; Dickerson et al., 2020; Buda et al., 2025). Moonlight has been associated with increased nocturnal vocal activity in some species and bird communities (Dickerson et al., 2020; Buda et al., 2025). However, natural and artificial light may not elicit equivalent behavioural responses. For example, experimental street lighting reduced nocturnal song in willie wagtails, with song activity recovering after the lights were removed (Dickerson et al., 2022). However, little is known about how different species within the same bird community respond to brief, discrete ALAN exposure and whether changes in nocturnal vocal activity are restricted to the illumination period or persist after the light source is removed.

In this study, we introduced a local artificial light source during nautical night and quantified vocalisation rates before, during, and after illumination. We hypothesised that vocalisation rates would increase during illumination relative to the pre-illumination phase and remain elevated after the lights were switched off. We also tested whether the magnitude and direction of the response differed among species and between early- and late-breeding-season surveys.

## METHODS

Experiments were conducted at ten sampling points in the Warta Landscape Park, central Poland, during nautical night, when the Sun’s geometric centre was more than 12° below the horizon. Experimental sites were separated by at least 200 m. We selected these sites for their high habitat diversity and lack of light pollution. Each sampling point was exposed to artificial light twice during the 2024 breeding season: once during the early survey, between 29 April and 4 May, and once during the late survey, between 6 and 14 June. Experiments were conducted under favourable weather conditions, with no rain or strong wind.

Five LED lamps were mounted on a stand 4 m above the ground and directed radially to illuminate the surrounding area. The experiment comprised four consecutive phases: 30 min under ambient darkness (before phase); an 8 min ramp-up period, during which one lamp was switched on every 2 min; 30 min with all lamps on; and 30 min after all lamps were switched off (after phase). The ramp-up period and the subsequent 30-min period of full illumination were combined into a single 38-min during phase.

AudioMoth 1.1.0 autonomous acoustic recorders continuously recorded bird vocalisations throughout the experiment. Vocalisations were identified to species by one observer (KB) using combined visual and auditory inspection in Raven Pro 1.6.5. The Supplementary Material provides further information on the experimental design, recording settings, light characteristics, environmental conditions, and vocalisation identification.

For each species, vocalisations were summed by sampling point, survey period, and experimental phase. We fitted negative-binomial generalised linear mixed models with vocalisation count as the response variable; experimental phase, survey, and their interaction as fixed effects; sampling point as a random intercept; and log-transformed recording duration as an offset. The offset accounted for the difference in duration between the 38-min during phase and the two 30-min phases. Formal analyses were restricted to species meeting the prespecified replication criterion. Species with insufficient spatial replication or unstable model estimates were described using raw observations. Detailed information on sample sizes, model structures, model diagnostics, false-discovery-rate correction, and pairwise comparisons are provided in the Supplementary Material and accompanying R script: https://github.com/kinkul1/Short-term-effects-artificial-light-at-night-on-nocturnal-vocal-birds-activity.git.

## RESULTS

We recorded 28,839 vocalisations from 13 bird species (Table 1; Fig. 1). Experimental phase significantly affected Eurasian coot vocalisation rate (χ^2^ = 12.15, pFDR = 0.011). The estimated rate was higher after illumination than before illumination (z = 2.88, p = 0.011) or during illumination (z = 2.56, p = 0.028), whereas the before and during phases did not differ (z = 0.51, p = 0.867; Fig. 1).

**Table 1.** Vocal activity recorded during the experiment. For each species and survey, the table presents the total number of vocalisations and the number of sampling points with at least one detection. Species used in formal statistical analyses are shown in bold.

| Species | Number of vocalisations |  | Number of sampling points |  |
| --- | --- | --- | --- | --- |
|  | Early survey | Late survey | Early survey | Late survey |
| **Common quail** | 970 | 16065 | 2 | 6 |
| **Sedge warbler** | 3232 | 2815 | 5 | 7 |
| **Common nightingale** | 4208 | 344 | 7 | 1 |
| Corn crake | 6 | 366 | 1 | 1 |
| Great reed warbler | 249 | 0 | 1 | 0 |
| Common cuckoo | 207 | 0 | 2 | 0 |
| **Eurasian coot** | 0 | 129 | 0 | 3 |
| Eurasian skylark | 116 | 0 | 3 | 0 |
| **Savi's warbler** | 37 | 49 | 1 | 3 |
| Whinchat | 0 | 39 | 0 | 1 |
| Eurasian bittern | 3 | 0 | 1 | 0 |
| Common crane | 2 | 0 | 1 | 0 |
| Mallard | 2 | 0 | 1 | 0 |

**Figure 1.**
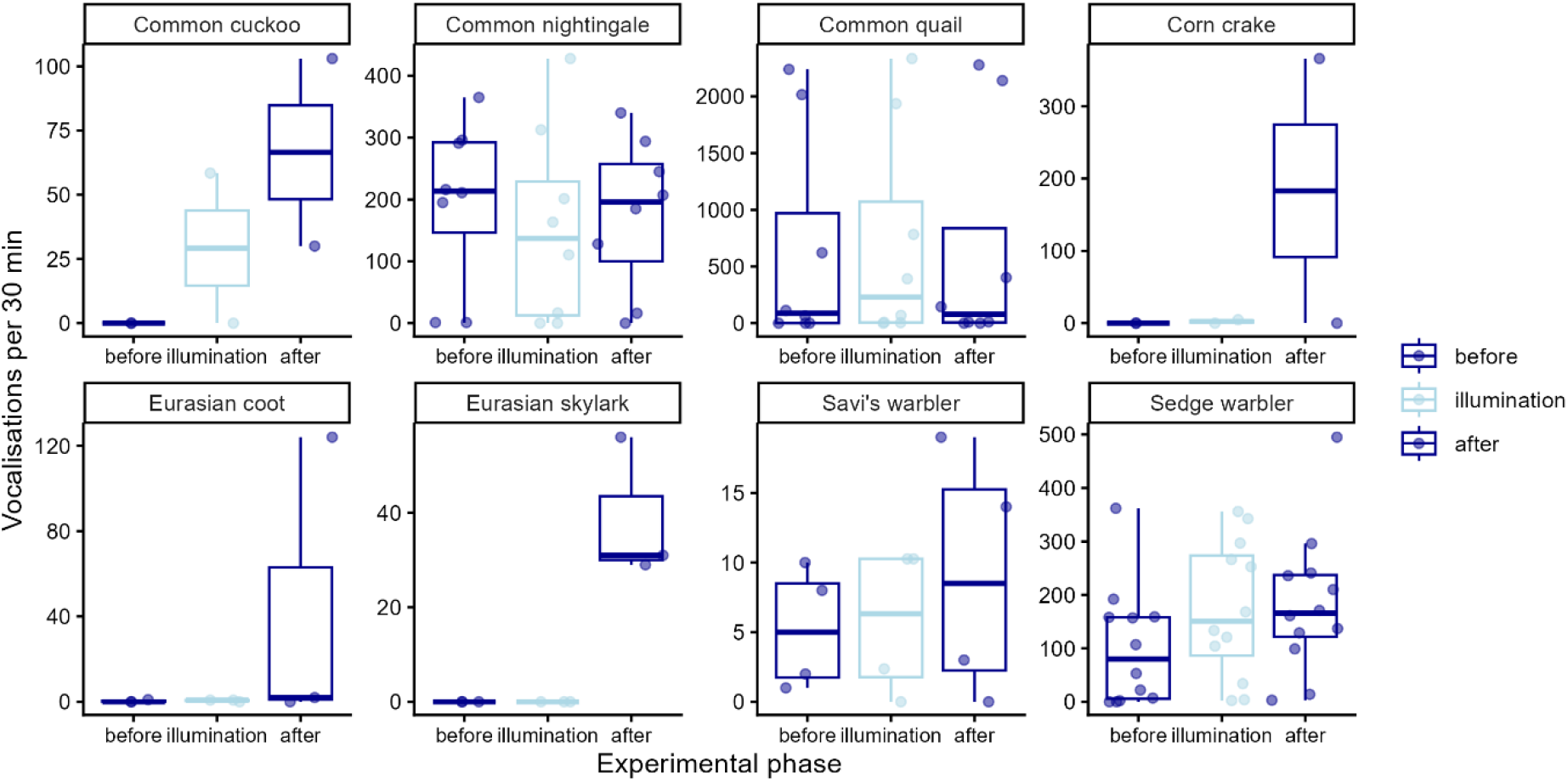
Vocalisation counts across experimental phases for species detected at two or more sampling points. Each point represents the number of vocalisations recorded at one sampling point during a single survey period, standardised to a 30-min sampling duration to account for differences in phase length: before = 30 min, illumination = 38 min, and after = 30 min. Boxes indicate medians and interquartile ranges, and whiskers extend to 1.5 times the interquartile range.

For sedge warbler, the overall effect of experimental phase was not significant after false-discovery-rate correction (χ^2^ = 6.45, pFDR = 0.099). Exploratory pairwise contrasts suggested higher song rates during than before illumination (z = 3.32, p = 0.003) and after than before illumination (z = 2.58, P = 0.027). Neither survey period (χ^2^ = 0.98, p = 0.322) nor the phase-by-survey period interaction (χ^2^ = 1.93, p = 0.382) effects were supported.

Eurasian skylarks were recorded at three sampling points only after illumination, with a median of 31 songs per 30 min (Fig. 1). This complete separation among phases prevented reliable model estimation. Consequently, the observed pattern is descriptive. Weaker patterns for common cuckoo and corn crake were also descriptive because each species was recorded at only two sampling points. Common cuckoo vocalisations were not detected before illumination but were recorded during and after illumination, whereas corn crakes were recorded only during the post-illumination phase.

We found no evidence of an experimental phase effect in common nightingale (χ^2^ = 3.55, PFDR = 0.283), common quail (χ^2^ = 0.22, pFDR = 0.894), or Savi’s warbler (χ^2^ = 0.59, pFDR = 0.894). Common quail vocalisation rate was higher during the late than during the early survey period (χ^2^ = 3.91, p = 0.048), but the phase-by-survey-period interaction was not supported (χ^2^ = 0.76, p = 0.685).

## DISCUSSION

Brief experimental illumination was associated with species-specific changes in nocturnal vocal activity. Of the five species analysed formally, only the Eurasian coot vocalised significantly more frequently after illumination than before or during illumination. The sedge warbler showed a tendency towards higher vocalisation rates during and after illumination than before illumination, although the overall effect of experimental phase was not statistically significant after correction for multiple testing. No significant differences in vocalisation rate among experimental phases were detected in the common nightingale, common quail, or Savi’s warbler. Descriptive patterns in three species not included in the formal analyses also suggested increased vocal activity: the common cuckoo vocalised more frequently during and after illumination, whereas the corn crake and Eurasian skylark showed an increase after illumination. Since the Eurasian skylark response is consistent for all 3 sampling points, the common cuckoo and corn crake response is not so clear, and should therefore be interpreted cautiously. Although previous morning point counts in Warta Landscape Park indicated that an average of approximately 7 bird species occurred within 100 m of the recording sites, only a small subset of the local bird community vocalised during the experiment (unpublished data; 25 points surveyed in 2024, with 4-15 species detected during individual 10-min surveys). This suggests that many species may show little or no immediate response to brief exposure to a local artificial light source, as no clear and consistent pattern was detected across the species recorded, at least over the short term. However, the present results are insufficient to determine the magnitude of ALAN effects at the community level. Greater spatial and temporal replication is needed to deeper understand community-level responses.

Previous studies show that ALAN can extend the daily period of avian vocal activity by advancing its onset around dawn and delaying its cessation around dusk, although the magnitude and direction of these effects vary among species and seasons (Pease & Gilbert, 2025). However, most studies have been correlational and could not fully separate the effects of ALAN from those of potentially confounding factors, such as noise pollution, urbanisation, or local environmental conditions. Experimental studies in which naturally dark habitats were illuminated throughout the night have produced contrasting results. Artificial illumination advanced dawn-song onset by approximately 4-37 min, depending on the species, in one study (Da Silva et al., 2017), whereas another found no evidence of earlier dawn singing in any of the 14 species examined (Da Silva et al., 2017). In addition, experimental ALAN reduced nocturnal song rates in willie wagtails, with song activity recovering after the light source was removed (Dickerson et al., 2022). Unlike these studies, our experiment examined short-term changes in vocalisation rates during and immediately after a brief illumination event in the middle of the night, rather than changes in dawn-song onset following all-night exposure. Our findings do not contradict previous evidence that ALAN can alter nocturnal vocal behaviour. Instead, they indicate that brief exposure may not elicit an immediate increase in vocalisation across species and identify candidate species that may show delayed or otherwise species-specific responses. Further controlled experiments are needed to determine whether ALAN exposure alters nocturnal vocal activity and whether any effects persist after the light source is removed

## Notes

### Competing Interest Statement

The authors have declared no competing interest.

### Funding

- Ministry of Science and Higher Education: This work was supported by the Diamond Grant programme of the Polish Ministry of Science and Higher Education, grant no. 0128/DIA/2019/48, “Nocturnal singing by diurnal birds – side effect of light or functional behaviour,” awarded to Kinga Buda.

