## Supplements for "Short-term effects of experimental artificial light at night on nocturnal vocal activity in a temperate bird community"

### Experimental details

During the illumination phase, we used 50 W PHOTON RGB LED floodlights as a local artificial light source. The lamps produced RGB light with a 120° beam angle and a rated power factor of 0.9. Five lamps powered by portable power banks were mounted on a stand 4 m above the ground and directed radially to illuminate the surrounding area. The lamps were switched on sequentially during an 8-min ramp-up period, with one additional lamp switched on every 2 min, and all five lamps then remained on for 30 min. At the end of the illumination phase, all lamps were switched off simultaneously.

Vocalisations at each sampling point were recorded using two AudioMoth 1.1.0 autonomous acoustic recorders. The recorders saved mono WAV files at a sampling rate of 48 kHz and a bit depth of 16 bits. They were attached to trees or shrubs 2-5 m from the lamps and 1-2 m above the ground. All experimental devices were operated automatically, and no observers were present during recordings. This minimised immediate disturbance associated with human presence.

Vocal responses recorded by the autonomous sound recorder were analysed in Raven Pro 1.6.5 software (Cornell Lab of Ornithology, Ithaca, NY, USA). Spectrograms were generated using a 1024-sample Hamming window. Each vocalisation was identified to species by combined visual and auditory inspection by one observer (KB).

### Detailed statistical analysis

Raw vocal recordings were first summarised as song counts for each combination of sampling point, species, month, and experimental phase. Missing combinations, where no songs were recorded, were assigned a value of zero. Combinations where a species was completely absent from a given point and survey period across all experimental phases were excluded because these observations could reflect a lack of occurrence rather than a response to experimental manipulation.

To evaluate species-specific responses to artificial light, we fitted separate negative binomial (nbinom2) Generalised Linear Mixed Models with a log-link function for six species. Only species recorded at more than three sampling points were included in formal statistical analyses. Experimental phase (three-level factor: before, illumination, after) was included as the main predictor to test whether artificial light affected vocal activity. Sampling point identity was included as a random intercept to account for repeated measurements at the same locations. Survey period was included either as a random intercept or as an additional fixed effect/interacting term depending on species-specific data structure, to account for repeated seasonal sampling or, if possible, differences in breeding phenology. For species with sufficient observations in both survey periods, the interaction between experimental phase and survey period was included to test whether responses to artificial light differed between stages of the breeding season. Finally, recording duration for each experimental phase was included as a log-transformed offset.

Model assumptions were evaluated using residual diagnostics, including tests for dispersion and zero inflation, implemented using the *performance* package (Lüdecke et al., 2021). The significance of fixed effects was assessed using Type II Wald $\chi^2$ tests implemented in the *car* package (Fox & Weisberg, 2011). Estimated marginal means of Experimental Phase were calculated on the response scale using the *emmeans* package (Lenth & Piaskowski, 2017), and pairwise comparisons among experimental phases were performed using Tukey-adjusted contrasts. Because multiple species-level tests were conducted, p-values from the experimental phase tests were corrected for multiple comparisons using the Benjamini–Hochberg false discovery rate (FDR) procedure.

All analyses were performed in R version 4.5.2 (R Core Team, 2025). Models were fitted using the *glmmTMB* package (Brooks et al., 2017), with data manipulation performed using *dplyr* (Wickham et al., 2014) and *tidyr* (Wickham et al., 2025) and visualisation using *ggplot2* (Wickham, 2016).
